# The Design Principles of Discrete Turing Patterning Systems

**DOI:** 10.1101/2020.10.18.344135

**Authors:** Thomas Leyshon, Elisa Tonello, David Schnoerr, Heike Siebert, Michael P.H. Stumpf

## Abstract

The formation of spatial structures lies at the heart of developmental processes. However, many of the underlying gene regulatory and biochemical processes remain poorly understood. Turing patterns constitute a main candidate to explain such processes, but they appear sensitive to fluctuations and variations in kinetic parameters, raising the question of how they may be adopted and realised in naturally evolved systems. The vast majority of mathematical studies of Turing patterns have used continuous models specified in terms of partial differential equations. Here, we complement this work by studying Turing patterns using discrete cellular automata models. We perform a large-scale study on all possible two-species networks and find the same Turing pattern producing networks as in the continuous framework. In contrast to continuous models, however, we find these Turing pattern topologies to be substantially more robust to changes in the parameters of the model. We also find that diffusion-driven instabilities are substantially weaker predictors for Turing patterns in the discrete modelling framework in comparison to the continuous case, and show that a more refined criterion constitutes a stronger predictor. The similarity of the results for the two modelling frameworks suggests a deeper underlying principle of Turing mechanisms in nature. Together with the larger robustness in the discrete case this suggests that Turing patterns may be more robust than previously thought.

## 1 Introduction

Nature is full of highly structured multi-cellular organisms that develop from single fertilized cells. It remains a key question how these can evolve robustly in the presence of environmental fluctuations.

The Turing mechanism has been proposed to explain such developmental patterning processes. It was first proposed by Alan Turing in 1952 [1]. The Turing mechanism gives rise to self-organised patterns in the local concentrations of biochemical components in reaction-diffusion systems. These lead to patterns such as spots, stripes and labyrinths [2]. Such inhomogenous patterns are induced by diffusion of the components. Due to this counter-intuitive concept, and the observation that Turing patterns are highly sensitive to initial conditions and variations in kinetic parameter values, the Turing mechanisms had been dismissed from the developmental community for almost two decades [3].

It was not until 1972 that Turing’s idea was revived by Gierer and Meinhardt who extended and formalised Turing’s work [4]. Despite some indications of suitable reaction-diffusion systems [5], the experimental technology available at the time was not able to convincingly identify Turing mechanisms in biological systems. It was not until another three decades later that technological advances have enabled compelling experimental evidence of Turing-like mechanisms [6]. Examples include the patterning of palatel ridges and digits, hair follicle distribution, and the patterns on the skins of animals, such as fish and zebras [7–11]. However, due to the complexity of the underlying systems, the exact molecular mechanisms are hard to identify most of the time, making it difficult to prove that the Turing mechanism exists in nature. The sensitivity to parameters constitutes another problem as it is not clear how biological systems could have evolved to find these small parameter ranges, and how these developmental processes can be so robust to extrinsic fluctuations.

This has resulted in a large number of theoretical studies in recent years [12–19]. Some recent studies have performed extensive analyses of potential network topologies [12, 14, 20]. Together these studies provide an inventory of the types of network structures that are capable of generating patterns and their robustness. The majority of these studies are preformed within a deterministic continuous framework in terms of partial differential equations (PDEs). An important question is if these results generalise to other types of modelling frameworks. This would indicate a deeper underlying principle of the Turing mechanism that is independent of the applied modelling framework. Moreover, since every model is an abstraction of a true biological system, the generalisation to different modelling frameworks would also suggest a certain robustness of the patterning processes.

Lattice gas cellular automata (LGCA) models constitute an alternative modelling framework [21]. These models discretize both space and the concentration of chemical components, and the dynamics is modelled by means of discrete diffusion and reaction steps. Turing patterns in LGCA models have first been studied in [22]. So far, only a single LGCA reaction map has been studied in this context. LGCA models have not been compared to continuous models in a broader scope with respect to Turing patterns (see also [23]). But they are ideally suited to studying developmental processes, where the cellular structure of tissues is explicitly included [24].

Here, we perform an exhaustive analysis of all possible two-species networks, and compare the results to the continuous modelling case. In our mathematical analysis of LGCAs we follow closely [22]. First, we analyse the network topologies with respect to diffusion-driven instabilities and analyse the emergence of patterns in simulations. In continuous models it was found that a diffusion-driven instability is a strong indicator of a pattern to emerge in simulations [20]. For LGCA models it has been shown for some specific systems that a diffusion-driven instability can also give rise to patterns in simulations. However, this is not always the case [22, 23]. It remains unclear how strong of an indicator a diffusion-driven instability is for pattern formation in the LGCA framework. Here, we analyse this exhaustively for all two-species models and show that a more refined instability criterion for LGCA models constitutes a better predictor for patterns emerging in simulations.

This article is structured as follows. In Section 2 we introduce the mathematical description of the LGCA, its simulation procedure and the definition of diffusion-driven instabilities. Next, we describe simulation details and how to identify patterns in simulations. Subsequently, in Section 3 we present the results obtained from our analysis of two-species networks and compare these results to the continuous case. Finally, we discuss the results and conclude in Section 4.

## 2 Methods

In this section we present the mathematical background on lattice gas cellular automata models (LGCAs) (Section 2.1), introduce the employed mean-field approximation (Section 2.3), define diffusion-driven instabilities (Section 2.3), and give simulation details in Section 2.4.

### 2.1 The LGCA model

We consider systems that consist of two interacting species modelled as discrete particles on a one-dimensional discrete lattice. The dynamics are modelled in discrete time steps that iteratively update the state of the system. Each update consists of separate reaction, shuffling, and diffusion steps, which are evaluated successively. Each spatial position consists of three compartments known as “velocity channels”, which can be either occupied or unoccupied by a single particle. Each spatial position hence can be occupied by a maximum of three particles (see supplementary material S1 for more details). Accordingly, we define the state of the system at time step *k* and lattice position *r* as

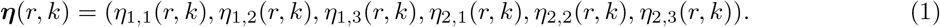

Each *η_i,σ_*(*r, k*) is a Boolean variable that represents the occupancy of velocity channel *i* of species *σ*, with *η_i,σ_*(*r, k*) = 1 (*η_i,σ_*(*r, k*) = 0) meaning the channel is occupied (unoccupied). Let further *n_σ_*(*r, k*) denote the total number of particles of species *σ* at spatial position *r* and time *k*,

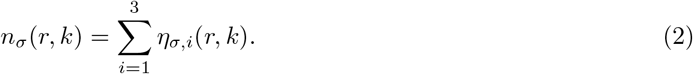

Next, let **X**= {0, 1, 2, 3} × {0, 1, 2, 3} denote the set of all possible states of (*n*_1_(*r, k*), *n*_2_(*r, k*)). We define the reaction step by a map **F**: **X**→ **X**, where each state **x** ∈ **X** is mapped onto **F**(**x**) = (*f*_1_(**x**), *f*_2_(**x**)) ∈ **X**. During the reaction step, each (*n*_1_(*r, k*), *n*_2_(*r, k*)) in each spatial position *r* is updated independently according to **F** with a probability 0 < *p* ≤ 1, and it remains the same with probability (1 − *p*). For *p* = 1 the reaction step becomes deterministic, and a smaller *p* introduces more stochasticity to the system. We refer to *p* as the “noise parameter”.

The *state transition graph* of the map **F** is a directed graph with the set of vertices **X**, and the set of edges (**x**, **F**(**x**))| **x** ∈ **X** for all **x** ∈ **X**. See Appendix A for more details.

The *reaction topology* of the map **F**, also known as “interaction graph” in the literature [25], is a graph that summarises the interactions between species: each species is represented by a node and each interaction is represented by an edge. Each edge is assigned a positive (negative) sign if the source node of the edge acts as an activator (inhibitor) on the target node. The edges can be identified from the state transition graph as follows: for species *i, j* ∈ {1, 2} there is a positive (negative) edge from species *j* to *i* if there exists a state **x** = (*x*_1_, *x*_2_) ∈ **X** such that *f_i_*(*x_i_, x_j_* + 1) − *f_i_*(*x_i_, x_j_*) is positive (negative). Intuitively, this means that an increase in *x_j_* is likely to lead to an increase (decrease) in *x_i_* (see supplementary material S2 for more details).

Note that due to the discrete nature of the state space there exist maps **F**: **X** → **X**, for which there exist **x**, **x**′ ∈ **X** such that *f_i_*(*x_i_, x_j_* + 1) − *f_i_*(*x_i_, x_j_*) > 0 and 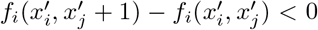, which means no single sign can be assigned to the edge from *j* to *i*. We omit such maps from our analysis since we are only interested in maps that can be assigned to the topologies in Figure 2.

**Figure 1:**
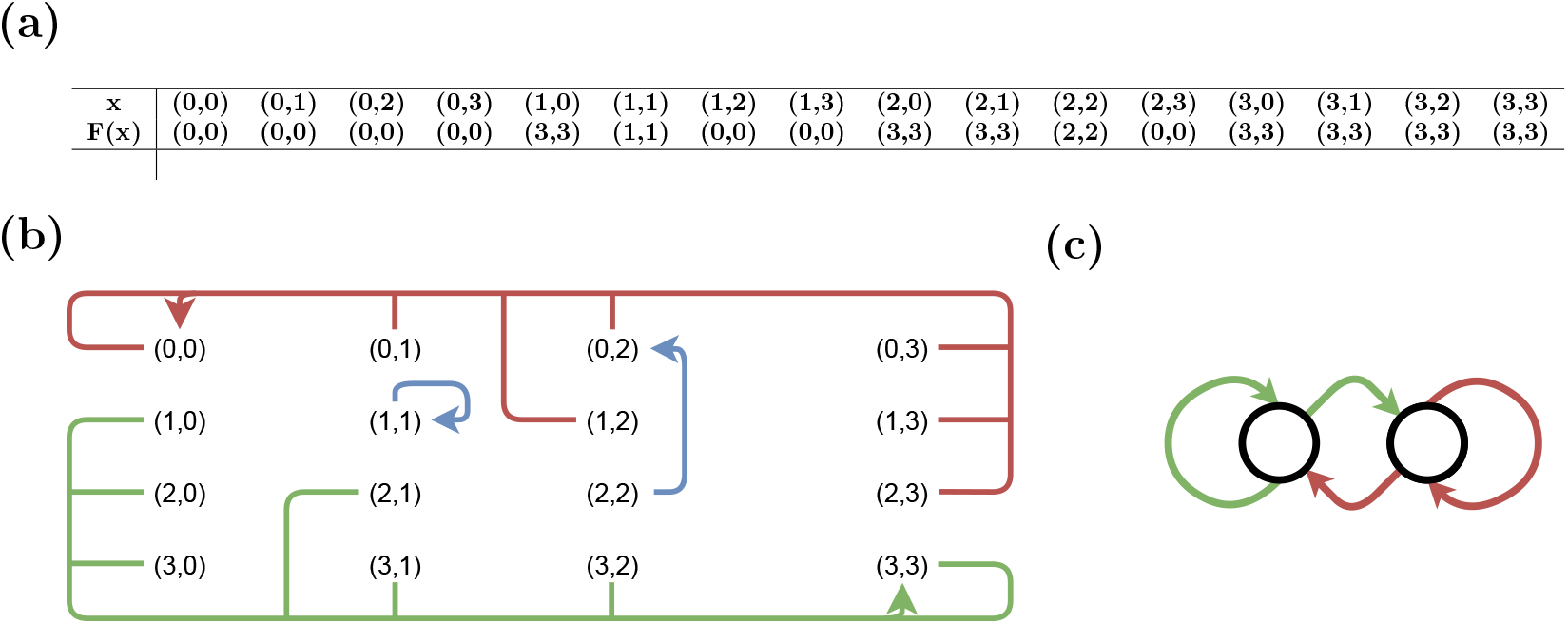
(a) The figure shows an example map **F** which determines the reaction step of the LGCA framework, i.e., it is a map from states **X** = {0, 1, 2, 3} × {0, 1, 2, 3} to states **X** = {0, 1, 2, 3} × {0, 1, 2, 3} and encodes the interactions of the species as described in Section 2.1, (b) the corresponding state transition graph and (c) the resulting reaction topology that summarises the interactions between the two species.

**Figure 2:**
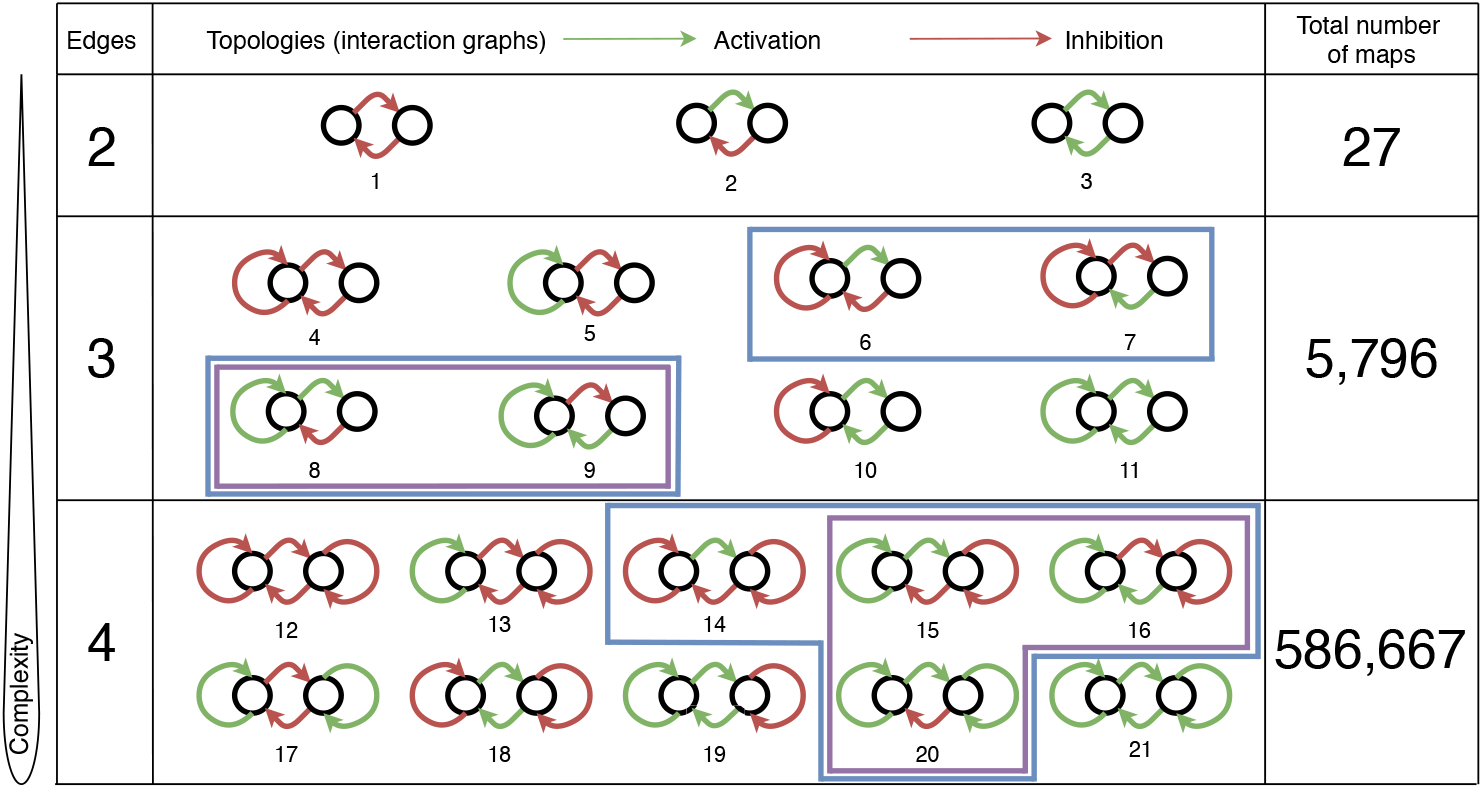
The 21 two-species reaction topologies distributed across three levels of complexity, where complexity is defined by the number of edges which corresponds to the number of interactions between the species. Highlighted in blue are the topologies that we find to contain maps producing diffusion-driven instabilities within the LGCA framework as defined in (I). Highlighted in purple are the topologies found to produce Turing instabilities within the PDE framework. We find these five to be the only topologies to produce patterns in LGCA simulations, as well as to be the exact same topologies that contain maps producing instabilities under the criterion in (II). The diffusion-driven instability criterion in contrast falsely predicts topologies 6, 7 and 14 to be able to produce patterns.

Figure 1 shows the function **F**, the state transition graph, and the reaction topology for the map originally analysed in [23].

Following the reaction step, the diffusion step acts simultaneously over all lattice positions and mimics a random walk of the particles on the lattice. It is comprised of two parts: a local random shuffling, where the particles of each lattice position are randomly redistributed across the three velocity channels; this is followed by a deterministic jump step, where a particle is moved on the lattice by a predetermined amount of spatial positions *d_σ_* ∈ ℕ_+_ in the direction associated with the velocity channel *i* it is occupying, *c_i_* ∈ {−1, 0, +1}.

We can formalise the diffusion step by defining the difference *C_σ,i_*(***η***(*r, k*), ***η***(*r* + *d_σ_c_i_, k* + 1)) of any given velocity channel *i* after one time step as

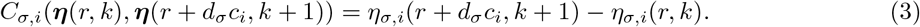

See appendix B for more details on the diffusion step.

### 2.2 Mean-field approximation

So far, we have defined the dynamics of the LGCA in an algorithmic manner which is convenient to simulate stochastic simulations but not convenient for mathematical analysis, in particular in the context of pattern formation. We consider the mean-field approximation as in [23], taking the expected value of Eq. (3) and neglecting correlations between channel occupations within a spatial position,

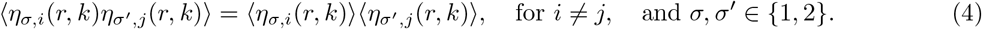

where 〈 · 〉 refers to the expectation with respect to the marginal distribution of *η_σ,i_*(*r, k*).

Using this one can derive the so-called “lattice-Boltzmann equations” which describe the evolution of the expectation values of the velocity channels [22],

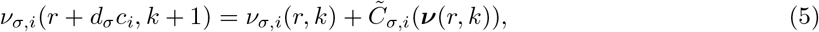

where *c_i_* denotes channel *i* direction of propagation, **c** = {−1, 0, 1}, ***ν***(*r, k*) = (*ν*_1,1_(*r, k*), …, *ν*_2,3_(*r, k*)), *C_σ,i_*(***ν***(*r, k*)) is now a function of time step *k* only (c.f. Appendix B, Equation (17)), and we define

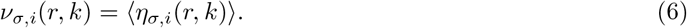

The *ν_σ,i_* are also referred to as “single-particle distributions” and can be viewed as the probability of finding a particle of species *σ* in channel *i* on lattice position *r* at discrete time point *k*. The stochastic shuffling of the particles across the channels within the diffusion step means that the single-particle distributions for each species *σ* ∈ {1, 2} are indistinguishable, so we define *ν_σ_* = *ν_σ,i_* for *i* ∈ {1, 2, 3} without loss of generality.

### 2.3 Stability analysis

We next define diffusion-driven instabilities in the context of LGCA using the lattice-Boltzmann equations in Eq.(5) by means of a linear stability analysis. A diffusion-driven instability describes the scenario where a stable steady state of a non-spatial system becomes unstable in the spatial setting where components are allowed to diffuse. By “stable” we mean that a small perturbation around the steady state asymptotically converges back to the steady state.

We start by defining a spatially homogeneous steady state 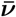 of the lattice-Boltzmann equations in Eq.(5) as the solution of

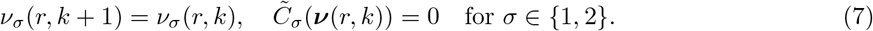

To assess the stability of a steady state 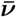 we analyse the evolution of a local perturbation of a certain wavenumber *q* around 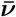 by applying a Fourier transformation to the lattice-Boltzmann equations. This decouples the individual frequency modes allowing the identification of unstable and stable modes. The evolution of the Fourier modes can be described in terms of the so-called “Boltzmann propagator” **Γ**(*q*) [26]. Here, **Γ**(*q*) has only two non-trivial eigenvalues *λ*_1_(*q*) and *λ*_2_(*q*) which can be derived analytically (see Appendix C for the derivation and expressions).

Analyses of the spectrum of the Boltzmann propagator aim at establishing the stability of each mode, and, as a consequence, the possible emergence of patterns [22, 23]. If |*λ*_1_(*q*)| < 1 and |*λ*_2_(*q*)| < 1 for all wave numbers *q*, one can expect the system to fluctuate around a stable spatially uniform solution, with no pattern emerging. In analogy with the continuous case, one could envisage the emergence of spatial patterns driven by diffusion if the spatially homogeneous steady state is stable but some modes can grow in time, that is, if the following conditions are verified:

I. |*λ*_1_(0)|, |*λ*_2_(0)| < 1 and |*λ_i_*(*q*\*)| > for some *q*\* > 0 and *i* = 1 or 2.

We refer to an instability that satisfies these conditions both as *instability type (I)* and *diffusion-driven instability*. We call the wave numbers *q*^∗^ for which *λ_i_*(*q*\*) becomes maximal the *dominant critical wave numbers*. [22] and [23] discuss some cases. They analyse examples where a critical wave number *q*\* ≠ 0 exists for which the corresponding dominant eigenvalue is real, and there is an indication of emerging spatial wavelength *L/q*\*. This is notably the case for an activator-inhibitor model that they study in detail, and whose reaction topology is given as number 15 in Figure 2. See Figure 3 (a) for an example simulation showing this behaviour and (b) for the corresponding eigenvalues as a function of *q*. Checkerboard structures seem to emerge if the dominant eigenvalue is smaller than −1, when oscillatory behaviours are expected (c.f. Eq. (23)). In contrast, for real eigenvalues larger than 1 stationary patterns seem more likely to emerge. They also identify examples of non-stationary spatial patterns, when the eigenvalue corresponding to a critical wave number has non-zero imaginary part. Since we are interested in stationary spatial patterns such as those emerging in activator-inhibitor models, for our analysis we consider the following criterion:

I. (II) the conditions in (I) are verified and *λ_i_*(*q*\*) corresponding to a dominant critical wave number *q*\* is real and positive.

**Figure 3:**
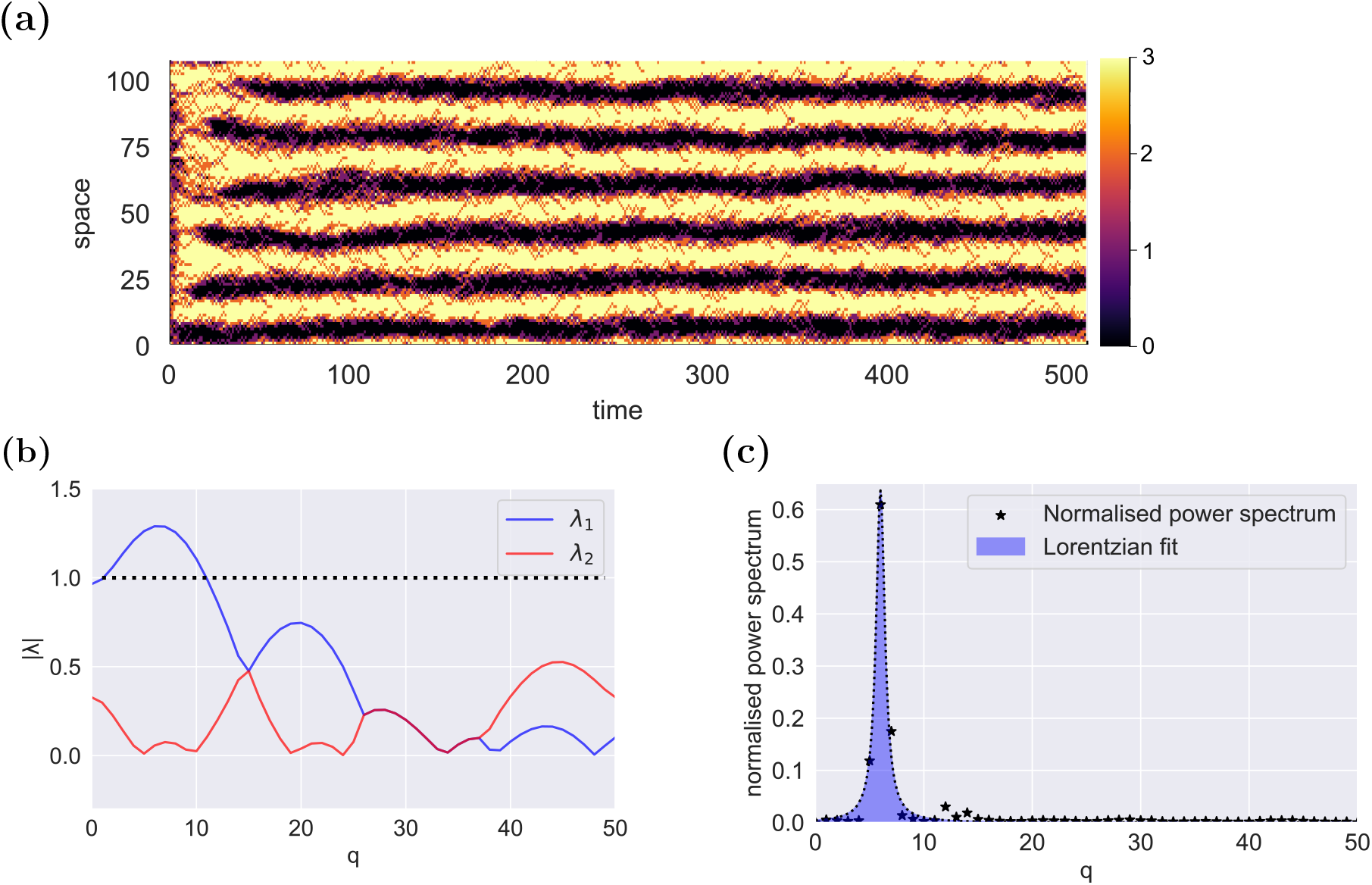
(a) Simulation of the map **F** shown in Figure 1. We find a clear pattern emerging. (b) Absolute values of corresponding eigenvalues *λ*_1_ and *λ*_2_ as a function of wave number *q*. The figure shows that the non-spatial system is stable, that is |*λ*_1_(0)|, |*λ*_2_(0)| < 1, while it becomes unstable for intermediate *q* values and maximal for a dominant wave number of *q*\* ≈ 7. This corresponds roughly to the wavelength *L/q*\* ≈ 14 observed in the pattern in (a). (c) Power spectrum of the simulation shown in b). The spectrum shows a significant signal at a frequency value of *q* ≈ 7, which corresponds to the dominant wave number *q*\* maximising the eigenvalue found in b).

We refer to an instability that satisfies these additional conditions as *instability type (II)*. For the two-species systems studied here we find that the dominant eigenvalues giving rise to diffusion-driven instabilities are always real-valued. The additional condition in (II) is hence equivalent to Re(*λ_i_*(*q*\*)) > 0 for the systems analysed here.

### 2.4 Simulation details and power spectrum analysis

Considering all different possible combinations of positive and negative edges between the two species gives rise to the 21 different fully-connected topologies shown in Figure 2 [20], i.e., topologies where both species influence each other. We only consider maps that can be assigned to one of these topologies. We thus exclude maps that have edges which cannot be assigned a positive or negative sign, as explained in Section 2.1.

We further reduce the number of maps to analyse by only considering asymptotic maps, which are defined by *f_σ_*(**x**) 0, *x_σ_*, ∈ {0, *x_σ_*, 3}, *σ* = 1, 2. This corresponds to a switch-like behaviour: the total occupation number of each species gets either updated to its maximal or minimal value, or remains the same. The total number of asymptotic maps for the 21 topologies of interest is 592, 490. Figure 2 shows the 21 possible reaction topologies grouped across three levels of complexity, which we define as the number of edges in the topology.

The formalism adopted here reduces the number of parameters considerably compared to the continuous case [20]. We set the noise parameter of the reaction step introduced in Section 2.1 to *p* = 0.9. We found empirically that *p* = 1 often leads to the system getting stuck in absorbing states that prevent the emergence of patterns; while smaller values of *p* introduce more stochasticity which tends to destroy patterns (see supplementary material S3 for examples).

Regarding the diffusion parameters *d*_1_ and *d*_2_ we found that the combination (*d*_1_, *d*_2_) = (1, 7) was the most likely combination to produce patterns for most studied reaction maps (results not shown). We also observed that small variations around this combination lead to negligible changes in the total number of maps that produce patterns. We hence fix (*d*_1_, *d*_2_) = (1, 7) throughout this article for simplicity.

To account for the stochasticity of the system, we simulate a given map a hundred times with random initial conditions in each simulation for *T* = 500 time steps on a domain of size *L* = 101 (see Section 2.1 for the simulation details). We then average each simulation over its last two time points to get rid of oscillatory spatial structures. Next, we compute the power spectrum via a Fourier transform, and average the result over the different simulation runs. Finally, we fit a Lorentz distribution to the maximum of the resulting average power spectrum (see Figure 3 (c) for an example and Appendix D and supplementary material S4 for details). The scale parameter *γ* of the fitted Lorentz distribution corresponds to the width of the peak at half its height, and is hence a measure for the peak width. A smaller *γ* value indicates a sharper peak in the Fourier transform and hence a clearer pattern in the simulation result. For the purpose of the analysis, we choose an arbitrary threshold (*γ* = 1) for *γ* to classify the maps as producing or not producing a pattern. The chosen threshold is close to the value giving the maximum F1 score (see Figure 7 (b)). The choice of a different threshold around this value does not impact the main conclusions of the analysis.

## 3 Results

In this section, we present the results obtained from both the linear stability analysis in Section 2.3, and stochastic simulations as outlined in Section 2.4. We start by discussing the results of the stability analysis before discussing different qualitative behaviours in simulations. We next present results on the emergence of patterns, and discuss different types of discrepancies between the stability analysis outcome and simulation outcomes. Finally, we discuss the robustness of the different topologies, and compare the LGCA results to the continuous modelling framework.

### 3.1 Stability analysis

We find that out of the 592, 490 asymptotic maps, 53, 479 possess a stable steady state, where steady states in the following are always defined in terms of the mean-field analysis as outlined in Section 2.2. We further find that a surprisingly large number of these^1^ (40, 519), which are more than 75%, possess a diffusion-driven instability (c.f. (I)) and 22, 965 of these fulfil the more refined instability criterion (c.f. (II)). We thus find that the refined criterion is substantially more restrictive, with it being fulfilled by only roughly half as many maps as the diffusion-driven one. Figure 4 visualises the hierarchical dependence of the different criteria, i.e. types of steady states, instabilities and simulation outcomes, together with the corresponding numbers of maps fulfilling them.

**Figure 4:**
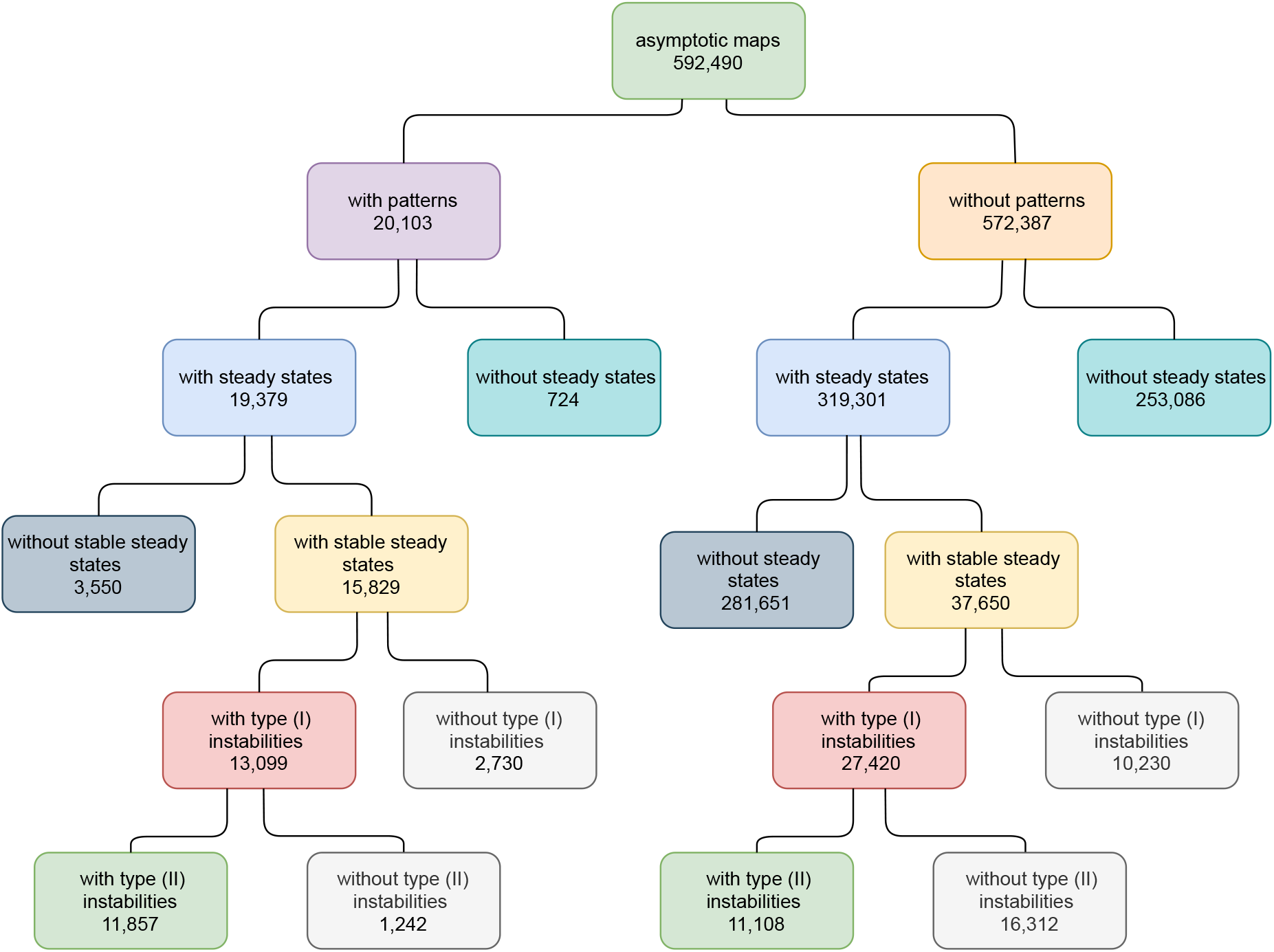
Illustration of the hierarchical dependence of the different analysed properties of maps for all asymptotic two-species maps. The corresponding numbers of maps fulfilling corresponding properties are also indicated.

The maps with diffusion-driven instabilities are distributed across eight reaction topologies (see Figure 2, highlighted in blue). All the maps within these topologies exhibit antagonistic behaviour between the two species, where the first species acts as an activator of the second species, while the second species inhibits the first. This activator-inhibitor principle is the generic mechanism used to describe Turing instabilities, first introduced by Gierer and Meinhardt [4]. Topology 8 corresponds to the classical Gierer-Meinhardt model (c.f. Figure 2), which consists of a slowly diffusing autocatalytic activator and a fast diffusing antagonist (inhibitor) species [4]. The other seven topologies with diffusion-driven instabilities are all variants of this core slow activator-fast inhibitor mechanism. Five of these eight topologies contain maps with instabilities that fulfill the conditions in (II). We refer to these topologies as “Turing topologies”.

### 3.2 Types of simulation outcomes

Diffusion-driven instabilities only describe the local instability of a steady state and do not guarantee the emergence of an actual pattern in simulations of a model. Indeed, in continuous PDE models it has been found that Turing instabilities do not always give rise to patterns in simulations [20]. Here, we find the same to be true for LGCA systems. We observe various different qualitative behaviours in simulations. Among these are oscillatory simulation outcomes which we disregard since we are interested in stationary patterns. We classify the remaining non-oscillatory outcomes into three categories: no structure emerging; structure emerging without a characteristic wavelength; and structure emerging with a characteristic wavelength (Turing-like patterns). We are interested in the latter: the “characteristic wavelength” of a pattern does not depend on the domain size (see Figure E).

Figure 5 (a) shows an example for the first case of no structure emerging, which means that either random noise emerges (as shown in Figure 5 (a)) or a spatially homogeneous state is reached. The corresponding power spectra do not contain significant signals indicating the absence of a pattern.

**Figure 5:**
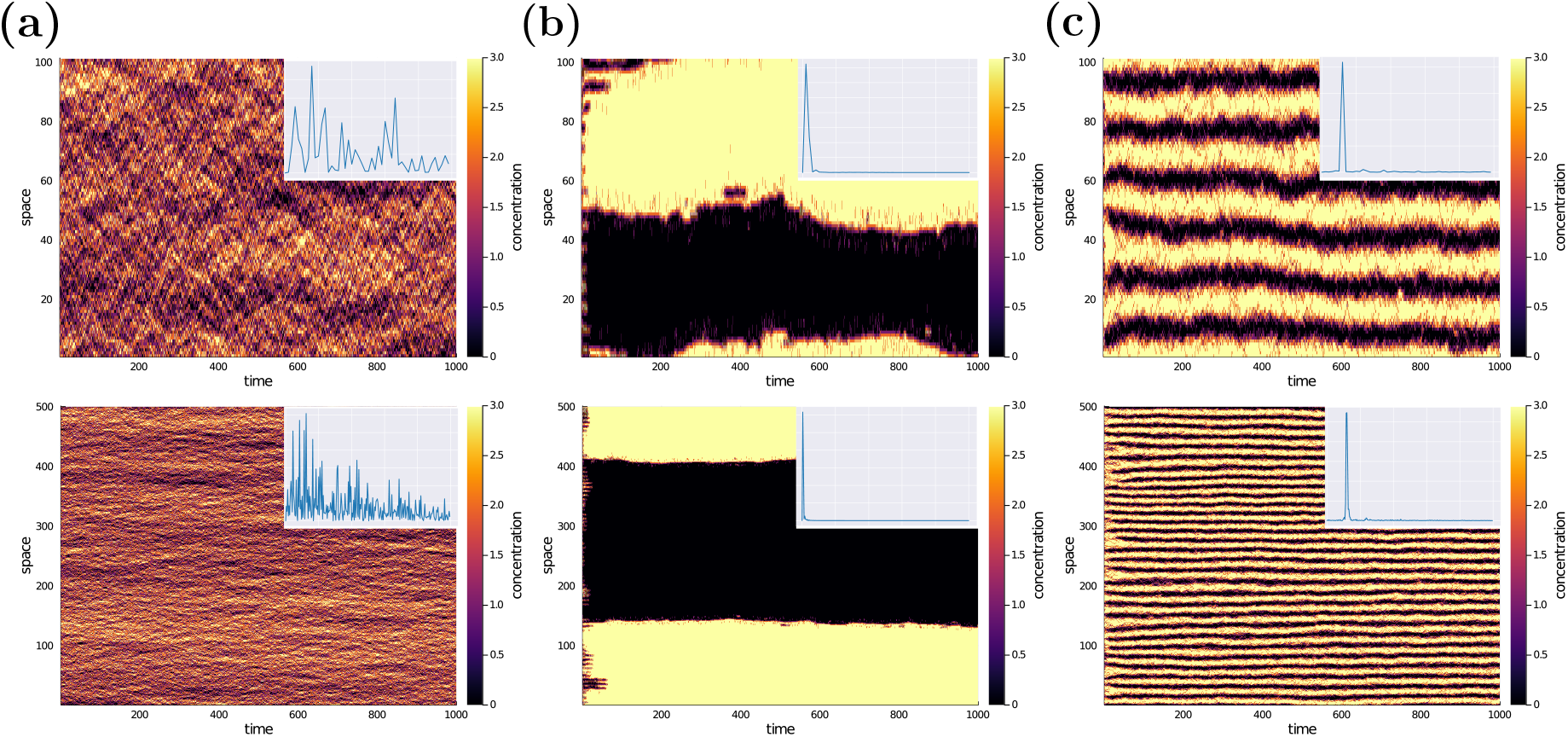
Examples of simulation outcomes for three different maps for two different domain sizes of *L* = 100 (top) and *L* = 500 (bottom) each. The insets show the corresponding power spectra (c.f. Section 2.4) (a) Example of no spatial structure forming. The corresponding power spectrum does not show a significant peak. (b) Example of spatial structure with no characteristic wavelength. The corresponding power spectrum shows a peak at a small wavelength of *q* = 1. (c) Example of spatial structure with a characteristic wavelength. The power spectrum shows a peak at an intermediate *q* which leads to a wavelength of *L/q* where *L* is the domain size. This is the type of pattern we are primarily interested in.

The second qualitative behaviour breaks the symmetry of the system and produces a structure without a characteristic wavelength. This behaviour is similar to the phenomenon of phase separation in PDE models [27]. The majority of these maps separates the domain into two parts, one highly expressed side and the other lowly expressed. Figure 5 (b) shows an example of a map that breaks symmetry with no characteristic wavelength. As the domain increases so does the resulting length-scale of the structure.

The third characteristic behaviour, to which Turing patterns belong, produces a stationary (i.e. non-oscillatory) spatial structure with a characteristic wavelength that does not depend on the domain size. Figure 5 (c) shows an example of this behaviour.

To automate the analysis of simulation results we proceed as explained in Section 2.4. For a simulation result to be classified as a pattern, we require the Lorentzian fitted to the normalised power spectrum averaged over 100 trials to have the scale *γ* ≤ *γ*\*, i.e., the peak must not be too broad. Note that this criteria is somewhat arbitrary, as the distribution of *γ* values for all maps is close to continuous. We found empirically that the threshold *γ*\* = 1 selects only maps that produce robust clear patterning..^3^

For some maps different simulation runs can give rise to different qualitative behaviours. For example, for some maps one observes simulations showing noisy patterns, and some simulations showing oscillatory behaviour or homogeneous solutions. The patterns give rise to clear peaks in Fourier space while the oscillatory or homogeneous solutions do not give such a peak. Averaging over several simulation runs will hence give an average *γ* value for the given map, which can be viewed as an effective measure of patterning capability. See supplementary material S4 for further details on this.

### 3.3 Instabilities as predictors for patterns

We find that only about 30% of the 40, 519 maps with diffusion-driven instabilities produce a pattern in simulations. Thus a diffusion-driven instability is only a weak predictor of a pattern to emerge. This is in stark contrast to the continuous case where only a very small fraction of systems with diffusion-driven instabilities have been found not to produce patterns [20]. For the 22, 965 maps fulfilling the refined instability criterion we find that about 50% lead to a pattern which suggests that this is a better predictor. However, there are also more maps giving rise to a pattern but do not fulfill this criterion. To make a more systematic comparison we hence consider the confusion matrices for each criterion, shown in Figure 6, which gives the numbers of maps categorised according to the four combinations of pattern/no pattern in simulations and instability/no instability, both for diffusion-driven and type (II) instabilities.

**Figure 6:**
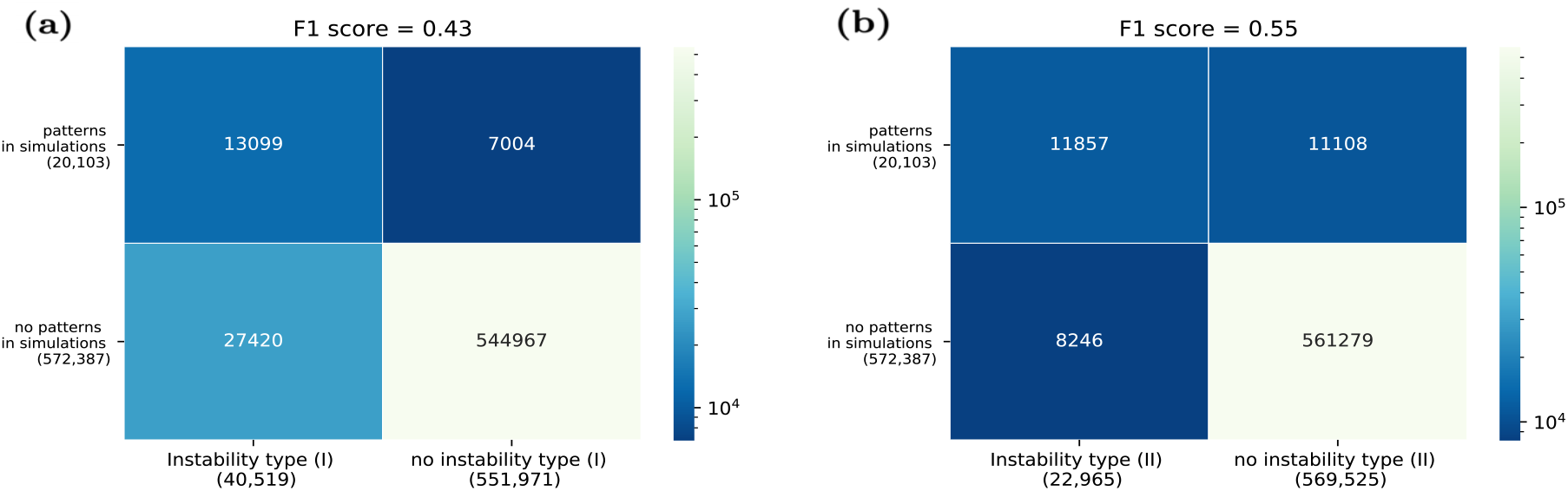
Confusion matrix for (a) the diffusion-driven instability criterion in (I); and (b) the confusion matrix for the instability criterion in (II). Shown are the numbers of maps that fulfill the respective indicated criteria. We find that the instability type (II) fails to predict patterns in simulations in only slightly fewer cases than the diffusion-driven instability (11, 857 vs. 13, 099 maps, see top left entries), while it predicts patterns falsely in substantial fewer cases (8, 246 vs. 27, 420 cases, see bottom left entries).

To quantify the overall predictability we note that the task of predicting patterns in simulations can be viewed as a binary classification problem, where the goal is to use the instability behaviour of a map as the input variable to predict the simulation outcome viewed as a binary target variable. We hence use the F1 score which is a standard measure to quantify the predictability for such binary classification tasks [28]. The F1 score is given by the harmonic mean of the precision and recall scores, where the precision is the ratio of correctly predicted patterns with respect to all maps with patterns, while the recall is the ratio of correctly predicted patterns with respect to all the predicted patterns [28]. A larger F1 score corresponds to a better predictability^3^. For diffusion-driven instabilities we find a F1 score of about 0.43 while for type (II) instabilities we obtain 0.55. We can therefore conclude that the instability criterion in (II) is a better predictor.

In terms of different topologies we find that the eight topologies 6, 7, 8, 9, 14, 15, 16 and 20 possess maps with diffusion-driven instabilities while only five of these, namely 8, 9, 15, 16 and 20, possess maps that give rise to patterns in simulations (see also Figure 2). The maps of topologies 6, 7 and 14 with diffusion-driven instabilities that fail to produce stationary patterns in simulations, mostly give rise to simulations that enter into spatially-homogeneous limit cycles, switching between high and low homogeneous expression levels (see supplementary S3 for examples). This is due to the eigenvalue giving rise to the instability being negative (c.f. Section 2.3), which was our motivation for the refined instability criterion. And indeed, we find that only topologies 8, 9, 15, 16 and 20 possess instabilities as in (II), giving a one-to-one correspondence between topologies giving rise to patterns in simulations and topologies possessing instabilities satisfying the more restrictive criterion (c.f. (II)). We will refer to these five topologies as “Turing topologies” in the following. We thus find the refined instability criterion to also be superior in terms of identifying the correct topologies.

So far we have used a threshold on the scale *γ* of the Lorentzian fit as a binary classifier to decide if a map produces a pattern or not. Instead, we can alternatively view *γ* as an inverse measure of the quality of a pattern: the smaller *γ*, the sharper the peak in the power spectrum. Figure 7 (a) shows the distribution of maps over the *γ* values for maps with diffusion-driven instabilities, for maps with type (II) instabilities, and for maps without instabilities. We find that the latter has a large peak at *γ* ≈ 15.9 which indicates a homogeneous simulation outcome, and only little weight for small *γ* values. The distributions of the two instabilities, by contrast, have large peaks for small *γ* values with substantial weight below the pattern threshold at *γ* = 1. The type (II) instabilities have more weight below this threshold and less above it in comparison to the diffusion-driven instabilities, indicating that they describe maps producing more distinct patterns on average.

**Figure 7:**
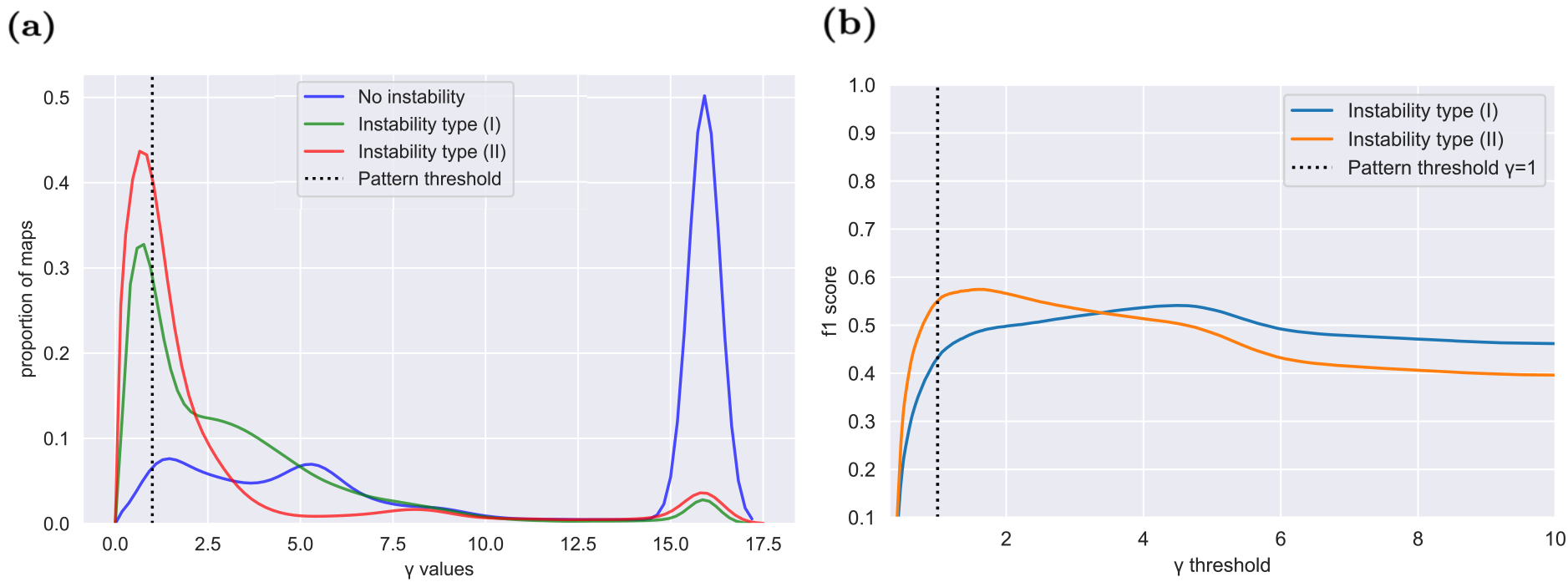
(a) Comparison of the *γ* distributions across sets of maps split into three categories: such with diffusion-driven instabilities (c.f. (I)); instabilities that satisfy the conditions in (II); and those without any instabilities. The dotted vertical line indicates the threshold *γ* = 1 below which we classify a simulation outcome as a pattern (c.f. Section 2.4). The peaks for large *γ* values around 15.9 stem from spatially homogeneous simulation outcomes. We observe that the distributions of maps with both instabilities have most of their mass at low gamma values, either close to or below the pattern threshold *γ* = 1. The ones without instabilities by contrast have only little weight for small *γ* values and a large peak beyond the signal threshold, i.e., the majority of maps gives rise to spatially homogeneous simulation outcomes. (b) F1 score of both instability types as a function of the chosen *γ* value for the pattern threshold. The dotted vertical line indicates the chosen threshold (*γ* = 1) for a pattern in our analysis. Instabilities of type (II) produce a better F1 score than instabilities of type (I) for all reasonable choices of pattern thresholds.

Since we find the instability type (II) criterion to be more meaningful than the diffusion-driven instability one in terms of predicting the existence of patterns, we will primarily focus on this in the following.

### 3.4 Discrepancies between stability analysis and simulations

As we have seen in Section 3.3, some maps with type (II) instabilities do not produce a pattern in simulations (the same is true for maps with diffusion-driven instabilities). This might be expected since the criterion only indicates a local diffusion-driven instability, and does not make statements about the global behaviour, as already mentioned in Section 3.2. Moreover, here we work with stochastic models, which means the random fluctuations can “wash out” the wavelengths emerging from an instability. Another reason for this discrepancy is that the system’s dynamics can get stuck in homogeneous absorbing states. The mean-field approximation used in the stability analysis may also contribute to such discrepancies. Another potentially important point is that we set a threshold for the scale parameter *γ* to define what we classify as a pattern.

As discussed in Section 3.3, we find 5, 285 maps that produce a stationary pattern despite not possessing a type (II) instability. These are all distributed across the Turing topologies, namely topologies 8, 9, 15, 16 and 20. 1, 011 of these 5, 285 maps possess a stable steady state, and we find that these maps all have an eigenvalue whose maximal absolute value is real, positive and close to one, and are hence somewhat “close” to a type (II) instability. One potential explanation why these give rise to patterns is that fluctuations present in the model can push the system over the threshold into the instability and hence lead to a pattern of the corresponding wavelength. This has also been suggested in [22]. The remaining 4, 274 maps do not possess a stable steady state, and the emergence of such patterns cannot be explained in the framework considered here, and therefore calls for further investigations.

### 3.5 Robustness

We next study the robustness of the different topologies [29]. By “robustness” we refer to the sensitivity of a topology’s pattern generation ability with respect to uncertainties in the model definition, i.e. with respect to the reaction map. We define this robustness as the fraction of maps of the topology that produces patterns in simulations. Figure 8 shows the fraction of maps that produce patterns for each topology. We find that for the Turing topologies a surprisingly large fraction of maps produce patterns, with fractions ranging from 0.032 to 0.28, meaning that up to 28% of asymptotic maps of some topologies are capable of producing stationary patterns. Interestingly, topology 8, which corresponds to the classical Gierer-Meinhardt model [4], was found to be the most robust topology by this measure. We also find that the robustness decreases with increasing complexity of the topologies (i.e. number of edges); this suggests that simpler models appear to be less sensitive to uncertainties in the precise model definition.

**Figure 8:**
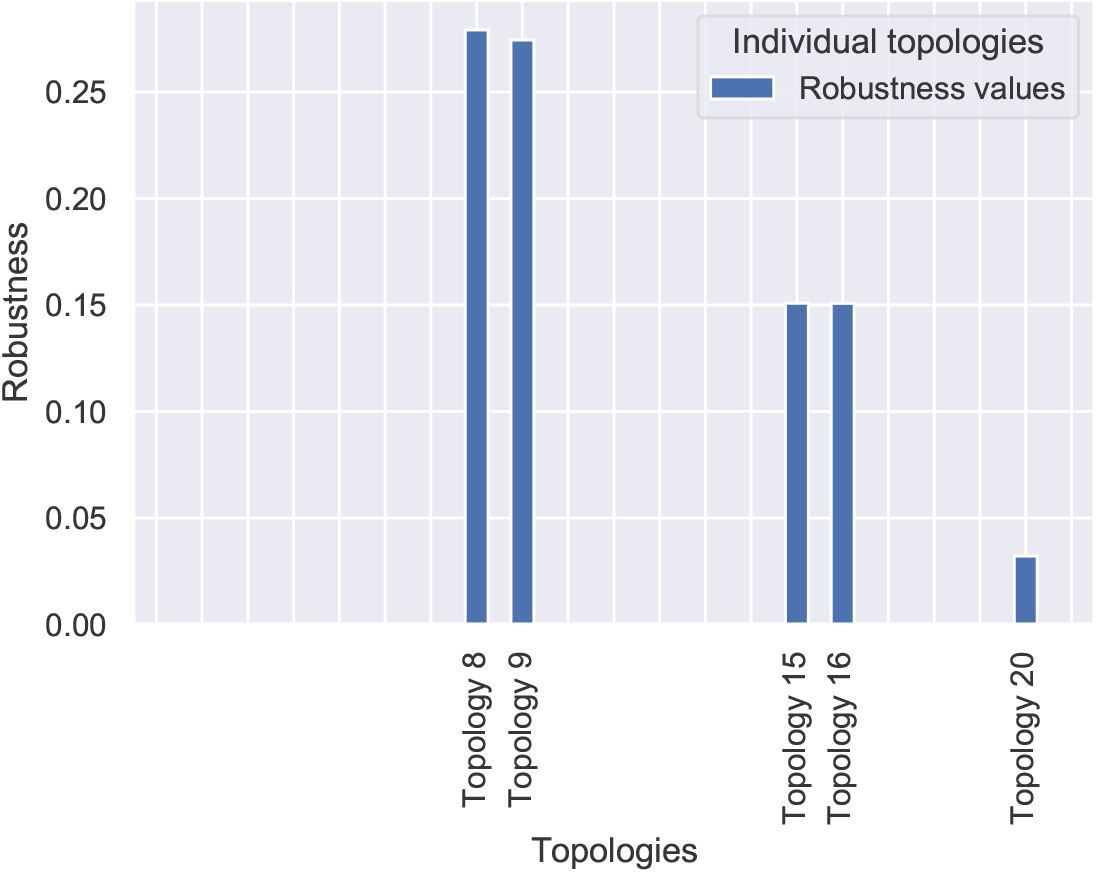
The figure shows the robustness values (defined as the fraction of maps producing patterns in simulations) for each topology. We find surprisingly large values of more than 28% for topologies 8 and 9. The robustness appears to decrease with increasing complexity (defined as the number of interactions between the two species, see Figure 2) to about 15% for topologies 15 and 16 and about 0.03% for topology 20. The other topologies have robustness 0 since they do not contain any maps that produce stationary patterns.

### 3.6 Comparison to continuous model

We next compare the results found so far using the LGCA approach with results from the literature using the continuous PDE approach. A direct comparison of individual systems of the two modelling approaches is not possible since we deal with discrete sets of reaction maps in the LGCA framework, while in the continuous case we have with continuously varying parameters. We thus compare the two approaches with respect to their overall behaviours in a broader sense [30].

#### 3.6.1 Instabilities and emerging patterns

In Figure 2 the five two-node topologies known to produce Turing instabilities in the continuous PDE case are highlighted in purple [12, 14, 20]. Here, we find that these five topologies produce type (II) instabilities and patterns in simulations in the LGCA framework. Therefore, in terms of patterns observed in simulations, we find a 1-to-1 correspondence between the topologies in the LGCA and the continuous frameworks.

#### 3.6.2 Instabilities as predictors of patterns

In Section 3.3 we found that 51% of the maps with type (II) instabilities produce a pattern in simulations (note that increasing the patterning threshold for *γ* would increase this number but would also increase the number of falsely predicted patterning maps, c.f. Section 3.3). This means that type (II) instabilities in the LGCA framework do not guarantee an emerging pattern in simulations. We found this to be independent of how many stable states a map possesses. In contrast, in continuous PDE models Turing systems with a single stable steady state have empirically been found to always give rise to patterns in simulations [20]. Only Turing systems with multiple steady states have been found to sometimes converge to homogeneous steady states in continuous models. Overall, type (II) instabilities that do not lead to patterns in simulations seem to be substantially more frequent in the LGCA framework. We may thus conclude that an instability in the LGCA formalism is a weaker predictor for the existence of a pattern than in the continuous case.

However, as already noted, the proportion of maps with type (II) instabilities that produce a pattern depend on our choice of the threshold for *γ*. Recall that *γ* can be viewed as a measure of how clear of a pattern one observes in simulations. In the continuous modelling framework, the amplitude of a pattern can be viewed as a similar measure of pattern quality. Some continuous models give rise to tiny amplitudes, and is questionable how relevant these are biologically and if they should really be considered as “patterns”. To the best of our knowledge this has not been considered in large-scale studies concerned with the robustness in continuous frameworks [12,14,20]. If one were to use a more sophisticated approach and apply a threshold on the amplitude of a simulation outcome in the continuous framework to classify them as patterns, the predictability of Turing instabilities would decrease accordingly. The disregard of pattern quality in continuous models should be taken into account when considering the relatively weak predictability of instability type (II) found here for the LGCA approach.

#### 3.6.3 Robustness

In Section 3.5 we defined the robustness of a topology in the LGCA framework as the fraction of maps that produces patterns in simulations. In the continuous modelling framework the robustness is typically defined as the fraction of kinetic parameters producing patterns in a given topology [12, 14, 20] (note that we do not consider robustness with respect to changes in diffusion constants or topology here as has for example been done in [20]). While being somewhat similar these two definitions are not equivalent and also depend on the considered set of maps and considered range of kinetic parameters, respectively. They can hence not be compared directly. Nevertheless, it is notable that the robustness values found here (ranging from 0.032 to 0.28) are substantially larger than values reported for continuous models (less than 0.01 for example reported in [20]). Applying an amplitude threshold as suggested in Section 3.6.2 would decrease the robustness found in continuous systems even further.

## 4 Discussion

Recent experimental findings [7–11] have resulted in Turing patterns being widely accepted as an important mechanism for spatial patterning in developmental processes. These findings have raised questions about the key features – or design principles – underlying the Turing mechanism and its robustness [29]. A variety of theoretical investigations aiming to answer these questions have been reported since. However, most of these have focused on single models [31–34]. More recently, some large-scale studies have systematically analysed large parts of possible design spaces, thereby providing a novel understanding on how common and robust Turing pattern mechanism are [12, 14, 20]. The majority of these theoretical studies have used differential equation models with continuous concentrations.

Since every mathematical model of a biological system is an abstraction, it is difficult to untangle those dynamics and features attributable to the true mechanics of a given biological system and those artificial dynamics arising from the modelling technique itself. Describing a biological system by different modelling frameworks can hence help to identify true underlying mechanisms; combining qualitative and quantitative modelling frameworks [35]; or augmenting modelling by evolutionary/comparative methods can provide reassurances as to the validity of modelling studies [36].

Here, we have performed a large-scale survey of all possible two-node topologies using a discrete lattice gas cellular automaton (LGCA) framework. This framework is much more restrictive than the continuous approach, in that it confines the concentrations to a small number of discrete values (four in our case) and comprises discrete maps between these states rather than continuous kinetic parameters.

Using this approach, however, we found the exact same five network topologies capable of producing Turing patterns as in the continuous modelling framework [12, 14, 20]. Moreover, we found these five topologies to be even more robust in the LGCA framework than they appear to be in the continuous case.

We also found that the presence of a type (II) instability, although a stronger predictor than a diffusion-driven (type (I)) instability, is neither sufficient nor necessary for pattern formation in the LGCA framework. Both criteria appear to be much weaker predictors of emerging patterns than Turing instabilities in the continuous counterpart. However, one has to keep in mind that studies assessing the predictability of instability criteria in continuous models typically do not assess the quality of the emerging patterns and therefore allow for arbitrarily small amplitudes. It is questionable how relevant patterns with arbitrarily small amplitudes are for noisy biological systems. Including a threshold on the amplitude in the definition of what constitutes a pattern in continuous models would presumably decrease the predictability of instabilities in these models, too. To some extent the smaller predictability of instability criteria in the LGCA framework might also be attributed to the stochasticity which makes the identification and definition of a pattern less straightforward, as stochasticity can wash out patterns and lead to breaking down of the employed mean-field analysis. With regards to network topologies, we found that, in contrast to the diffusion-driven instability criterion, the refined criterion (type (II)) identifies the same five topologies capable of producing patterns in simulations. Our results thus suggest that the type (II) instability is a better predictor of patterns in simulations than the diffusion-driven instability (type (I)).

The fact that the restricted model considered in this work identifies the same Turing topologies identified before in the continuous case suggests that the exact molecular details might not be as important as previously thought and hints at a deeper underlying principle of Turing mechanisms that is independent of the modelling framework. The larger robustness we found also suggests that Turing patterns might be more robust than previous continuous studies suggest.

### A Reaction step

Following from Section 2.1 we derive the general expression for the channel after the reaction step. Recall from Section 2.1 that the total number of particles of a given species *σ* at a given position *r* and time *k* is

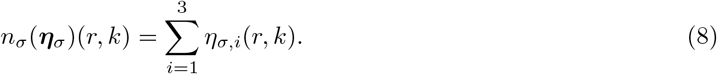

After the reaction step the newly updated total number of particles of species *σ* at a given *r* and time *k* is

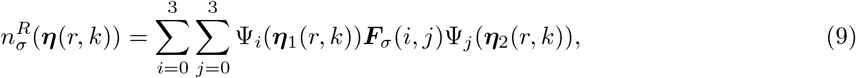

where the *indicator function* Ψ is defined as Ψ(***η***_*σ*_(*r, k*)) = (*ψ*^0^(***η***_*σ*_(*r, k*)), *ψ*^1^(***η***_*σ*_(*r, k*)), *ψ*^2^(***η***_*σ*_(*r, k*)), *ψ*^3^(***η***_*σ*_(*r, k*))) with *ψ^i^*(***η***_*σ*_(*r, k*)) = 1 when *i* particles of species *σ* exist at the lattice position *r* and 0 otherwise. See supplementary S5 for a full expression of Ψ(***η***_*σ*_(*r, k*)). **F**_*σ*_(*i, j*) denotes the updated value of species *σ* from the state **x**= (*i, j*) as defined in section 2.1. The superscript ^*R*^ indicates the variable after the reaction step. We add stochasticity to the interaction step by introducing sequences of space and time independent identically distributed Bernoulli random variables ϵ ∈ *E* {0, 1}, 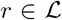, *k* ∈ ℕ. These variables determine whether the reaction takes place or not. We further define *p* = *P* (ϵ(*r, k*) = 1), where *p* is the probability of the reaction taking place. We refer to *p* as the “noise parameter”. With these definitions the post-reaction total number of particles of species *σ* at a given *r* and time *k* reads

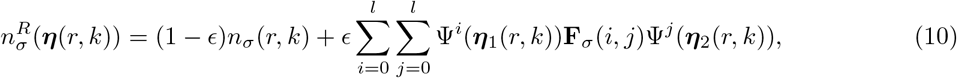

Once the total number of particles 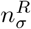 are updated they are redistributed back into the individual channels 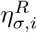 for all *i*, such that

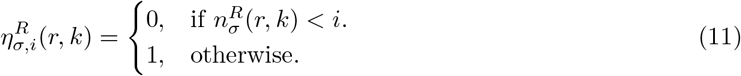

### B Shuffling step, diffusion step and difference equation

In this section we provide a formal definition of the shuffling and diffusion steps described in Section 2.1, and give the full expression for the difference equation in (3). We start by expressing the random shuffling step in terms of permutation matrices. The set of permutation matrices for a system with three velocity channels is

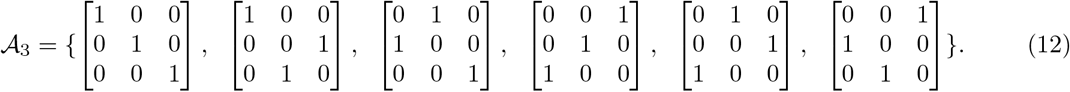

Using these we can write the updated channels in terms of the set of local channels at spatial position *r* as

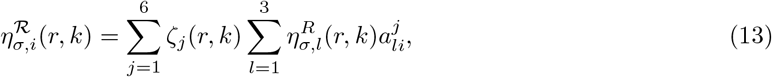

where the superscript 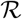 indicates the variable after the shuffling step. 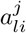 denotes the element of the *j^th^* permutation matrix at row *l* and column *i*, and *ζ_j_* ∈ {0, 1}, *j* ∈ 1, … 6 are Bernoulli type random variables, such that *ζ_j_* = 1 for one *j* ∈ {1, …, 6} and zero otherwise. After the shuffling step we apply the jump diffusion step as

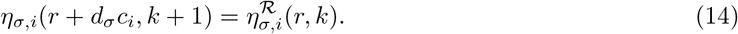

Combining Equations (3), (13) and (14) we obtain an expression for the change in expression level over a single time step, *C_σ,i_*(***η***(*r, k*), ***η***(*r* + *d_σ_, k* + 1)), in terms of the variables of the previous time step:

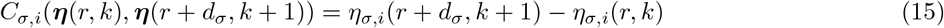

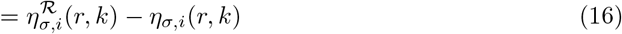

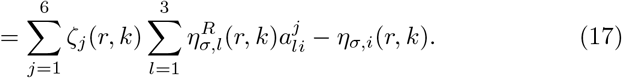

and we can write *C_σ,i_*(***η***(*r, k*), ***η***(*r* + *d_σ_, k* + 1)) = *C_σ,i_*(***η***(*r, k*)). Equation (17) describes how each individual channel evolves over time.

### C Linear stability analysis in the LGCA model

Here, we derive the equations needed for the stability analysis used to define the instability criteria in the mean-field approach in Section 2.3. As mentioned in Section 2.3, we use a small perturbation to determine the stability of a given steady state: let 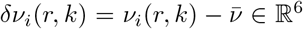 be a small perturbation around the steady-state solution 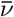 of Equation (7), where *i* denotes the channel. Using this in (5) and linearizing around 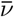 we get the linear lattice-Boltzmann equation

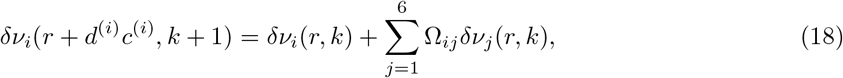

where the diffusion coefficient *d*^(*i*)^ is the *i*th element of (*d_σ_*_=1_, *d_σ_*_=1_, *d_σ_*_=1_, *d_σ_*_=2_, *d_σ_*_=2_, *d_σ_*_=2_) and the direction *c*^(*i*)^ is the *i*th element of (1, 0, −1, 1, 0, −1). The Jacobian **Ω**∈ ℝ^6×6^ is defined as

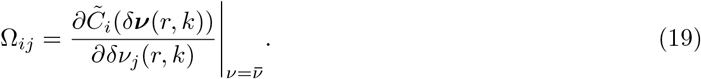

Consider a harmonic wave perturbation of the form

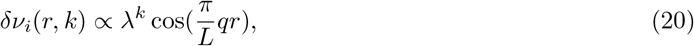

where *q* = 0 corresponds to a spatially homogeneous perturbation. Next, we consider a general perturbation **F**(*q, k*) = (*F*_1_(*q, k*), …, *F*_6_(*q, k*)) and express each of its components as a sum of sinusoidal terms as

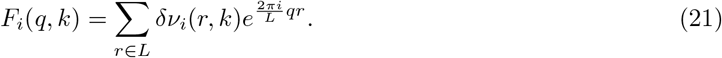

Applying the discrete Fourier transformation to the linear lattice-Boltzmann equations (18) gives

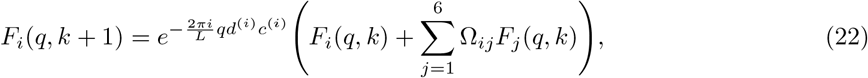

which we write in vectorised from for as

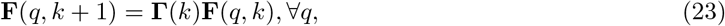

where **Γ**(*k*) is the Boltzmann propagator defined by

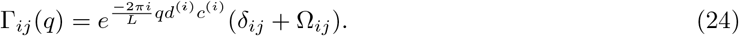

In matrix notation this can be written as

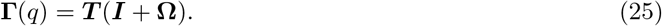

where **T**∈ ℝ^6×6^ which is known as the “Transport matrix” and defined as a diagonal matrix with elements 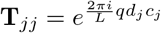, and **I**∈ ℝ^6×6^ is the identity matrix. (**I**+ **Ω**) is a block matrix and reads

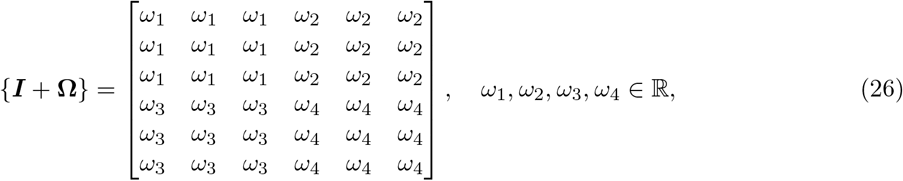

where the *ω_i_* are steady-state dependent constants and defined in the supplementary material S5.

We define **Λ**_Γ(*q*)_ = (*λ*_1_(*q*), *λ*_2_(*q*), *λ*_3_(*q*), *λ*_4_(*q*), *λ*_5_(*q*), *λ*_6_(*q*)), where *λ_i_*(*q*) is the *i*th eigenvalue of **Γ**(*q*).

The *λ_i_* determines the stability of a given steady state and are obtained as solutions of

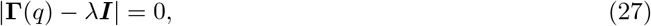

where | · | denotes the determinant. Due to the block structure of **Γ**(*q*) (c.f. equation 26) only two of the eigenvalues are non-zero: **Λ**_Γ(*q*)_ = (*λ*_1_(*q*), *λ*_2_(*q*), 0, 0, 0, 0), with

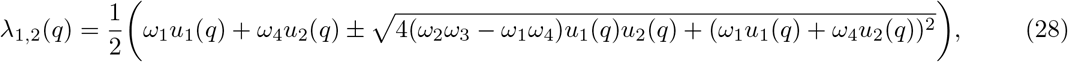

where 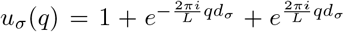. When *q* = 0, *λ*_1_ and *λ*_2_ define the stability of the non-spatial system.

### D Power spectrum analysis

In this section we provide details on how we use the power spectrum in Section 2.4 to automatically identify patterns in simulations as outlined in Section 2.4. Since either both or none of the two species show patterns in simulations, we only use the number of particles *n*_1_(*r, T*) and *n*_1_(*r, T* − 1) of the first species defined in Equation (2) at the last two time points *T* and *T* − 1 to identify patterns. The normalised power spectrum of a simulation run as described in Section 2.4 is given by

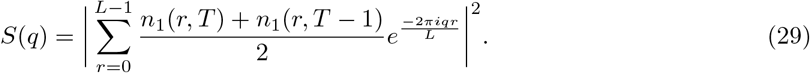

The average over the last two time points is taken to get rid of oscillatory spatial structures that would produce significant peaks in the power spectrum at single time points. To average out fluctuations we perform *K* simulation trials. Let *S*(*q*)^(*i*)^, *i* = 1, …, *K* be the corresponding power spectra computed as in Equation (29). We accordingly define 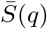 as the average over the *K* trials:

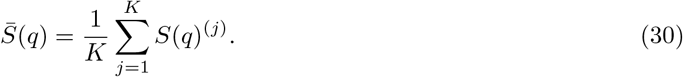

If 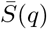 possesses a clear peak at a certain wavenumber *q* this indicates a spatial pattern in the simulation results with wavelength *L/q*. To identify such a peak we fit a Lorentz distribution to 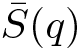 to quantify the quality of a pattern. The Lorentz distribution is defined as

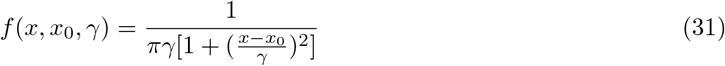

[37]. We determine the median *x*_0_ by finding the frequency in which the cumulative distribution function is equal to 0.5. We then use the method of least squares to determine the value of the scale parameter *γ* ∈ [0, 1]. The smaller the scale parameter the sharper the peak and hence the clearer the pattern in the spatial domain. We thus use a threshold on *γ* to judge if a simulation result contains a pattern or not, as explained in Section 2.4.

### E Identifying patterns with characteristic wavelengths

In Section 3.2 we discuss three types of simulation outcomes. Two of these correspond to spatial pattern-like structures that can arise: one with and one without characteristic wavelengths. A spatial structure is said to have a characteristic wavelength if the wavelength is independent of the spatial domain size. Figure 5 b) and c) show spatial structures produced over two different spatial domain sizes. Figure c) shows patterns with the same wavelength for both domain sizes, meaning the structure has a characteristic wavelength, whereas Figure b) shows patterns with the same bisection of the spatial domain into two halves. The wavelength therefore depends on the domain size and is not characteristic of the system that produced it.

Since we are interested in patterns with characteristic wavelengths we need to distinguish them from patterns that do not possess a characteristic wavelength. To this end we compare the wavelengths obtained by simulating a map on the two domain sizes *L* = 100 and *L* = 500. The wavelength for a given simulation is given by *L/q* where *L* is the domain size and *q* is the location of the peak in the power spectrum (c.f. Figure 5). If the two wavelengths obtained for the two domain sizes deviate by less than 10% from each other we consider them to be the same and conclude that the pattern has a characteristic wavelength.

## Footnotes

1 Recall that both type (I) and type (II) instabilities are only defined for maps with a stable steady state, c.f. Section 2.3

2 It should be noted that in the continuous case a similar problem arises: some systems with Turing instabilities give rise to patterns with tiny amplitudes. It is questionable how biologically relevant such systems are and it might be sensible to apply some cutoff here on the amplitude of the pattern too to define what an actual pattern is.

3 Note that the obtained numbers are dependent on the chosen threshold for *γ* in the definition of what is classified as a pattern (c.f. Section 2.4). If we choose a larger threshold for *γ*, then the ratio of maps with an instabilities that produce a pattern would increase. However, so would the number of maps that produce patterns but lack the instability. The F1 score takes into account this trade-off. Figure 7 (b) shows the F1 score as a function of this threshold choice, it shows that for all reasonable choices of threshold instabilities of type (II) produce a better F1 score than instabilities of type (I).

